# Confirmation bias is adaptive when coupled with efficient metacognition

**DOI:** 10.1101/2020.07.28.225029

**Authors:** Max Rollwage, Stephen M. Fleming

## Abstract

Biases in the consideration of evidence can reduce the chances of consensus between people with different viewpoints. While such altered information processing typically leads to detrimental performance in laboratory tasks, the ubiquitous nature of confirmation bias makes it unlikely that selective information processing is universally harmful. Here we suggest that confirmation bias is adaptive to the extent that agents have good metacognition, allowing them to downweight contradictory information when correct but still able to seek new information when they realise they are wrong. Using simulation-based modelling, we explore how the adaptiveness of holding a confirmation bias depends on such metacognitive insight. We find that the behavioural consequences of selective information processing are systematically affected by agents’ introspective abilities. Strikingly, we find that selective information processing can even improve decision-making when compared to unbiased evidence accumulation, as long as it is accompanied by good metacognition. These results further suggest that interventions which boost people’s metacognition might be efficient in alleviating the negative effects of selective information processing on issues such as political polarisation.

## Introduction

Polarization between opposing viewpoints is increasingly prevalent in discussions surrounding political and societal issues (Kohut et al., 2012). An important cognitive driver of this polarization is the human tendency to discount evidence against one’s current position (Del Vicario et al., 2017; Lilienfeld et al., 2009; Rollwage et al., 2018, 2019), a phenomenon known as confirmation bias (Nickerson, 1998). Confirmation bias has been reported in a variety of settings (Klayman, 1995), including the formation of clinical diagnosis (Groopman, 2008), inference about people’s character (Snyder & Swann, 1978), investment decisions (Park et al., 2010), views about societal issues such as capital punishment (Lord et al., 1979) and climate change (Sunstein et al., 2016). Perhaps most prominently, confirmation bias has been reported in relation to politically charged beliefs, such that people are generally prone to process information in line with their political convictions (Kaplan et al., 2016; Nyhan & Reifler, 2010; Redlawsk, 2002; Taber et al., 2009; Taber & Lodge, 2012). On a societal level, such skewed information intake might lead to entrenched beliefs, and, in turn, the prevalence of dogmatic groupings and widespread polarization (Zmigrod, 2020; Rollwage et al., 2019). In line with this hypothesis, people who show a resistance to belief updating are also more likely to show extreme political beliefs (Zmigrod, Rentfrow, et al., 2019b), aggression towards opposing political views (Zmigrod, Rentfrow, et al., 2019a), and authoritarian (Sinclair et al., 2019) or dogmatic traits (Rollwage et al., 2018; Zmigrod, Zmigrod, et al., 2019).

Recent cognitive neuroscience studies have identified the selective integration of choice-consistent information as a key mechanism underpinning this cognitive bias (Palminteri et al., 2016; Rollwage et al., 2020; Talluri et al., 2018). For instance, we recently showed that dogmatic participants were characterized by two cognitive alterations in the context of a perceptual decision-making task (Rollwage et al., 2018). First, dogmatic participants showed a reduction in metacognitive ability, manifesting as a selective overconfidence after making errors. Second, metacognitive ability (i.e. the accuracy of confidence judgments) was predictive of post-decision evidence integration, where people with poorer metacognition showed less sensitivity for corrective information.

These results indicate that confidence may act as an internal control signal that guides future information processing (Atiya et al., 2019; Balsdon et al., 2020; Desender et al., 2018; Meyniel & Dehaene, 2017; Murphy et al., 2015). In studies using magnetoencephalography (MEG), we found evidence for this hypothesis, with confidence strongly modulating the extent of neural post-decision processing. Evidence accumulation was largely unbiased after low confidence decisions but displayed a confirmation bias after high confidence decisions (Rollwage et al., 2020). In other words, people appear especially resistant to corrective information when they are highly confident about a wrong decision. However, when confidence is well aligned with performance – when metacognitive ability is high – weighting post-decisional integration by confidence is likely to be less problematic, as people will tend to be less confident after making errors, and therefore also open to corrective information. This line of reasoning suggests that people’s metacognitive ability might be a crucial driver of the degree to which selective information processing leads to negative behavioural outcomes.

Here we test a hypothesis that selective information integration might be adaptive when coupled with high metacognitive ability. Our proposal is in line with a broader hypothesis that the ubiquitous nature of selective evidence integration makes it unlikely that this cognitive characteristic is always maladaptive (Gigerenzer, 2008; Klayman & Ha, 1987). For instance, others have interpreted confirmation bias as a heuristic that reduces computational complexity (Evans, 1989) or allows for robustness against noise (Lefebvre et al., 2020; Tsetsos et al., 2016). Here we offer an alternative perspective: that confirmation bias is adaptive to the extent it is accompanied by a metacognitive ability to effectively monitor and recognise when we might be wrong (Fleming & Dolan, 2012). We use simulation-based modelling to compare different evidence integration strategies and test their respective performances. Specifically, we compared unbiased evidence integration with a simple confirmation bias as well as with a confidence-driven confirmation bias (as observed empirically in Rollwage et al., 2020). Since we expected the performance of a confidence-weighted confirmation bias to depend on the reliability with which confidence judgments indicate choice accuracy, we also investigated the influence of metacognitive ability on the adaptiveness of confirmation bias.

### Modelling behavioural impact of selective evidence integration strategies

We model a simple situation in which agents make a binary decision between two choice options based on noisy information drawn from a world state that is unknown to the agent. This situation is easily accommodated by existing frameworks for characterising belief updating (Bronfman et al., 2015; Fleming et al., 2018; Talluri et al., 2018), and closely resembles common perceptual decision-making paradigms. This setting also acts as a minimal framework within which more complex decision problems can be modelled. For instance, to reach an opinion about whether human activity causes global warming (the ground truth), we have to form beliefs based on multiple noisy evidence samples (e.g. scientific publications and newspaper articles). Importantly, this process requires the updating of pre-existing beliefs whenever more evidence becomes available. In such a situation, Bayesian belief updating is often used as benchmark model (Mathys et al., 2011; O’Reilly et al., 2013). Thus, we incorporated a Bayesian updater as our “unbiased agent” against which other evidence integration strategies can be compared. A Bayesian agent keeps track of its graded belief that one or other choice option is correct by calculating the probability of the chosen option, given the evidence, compared to the alternative: 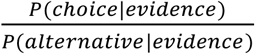

For simplicity, we simulate a situation in which participants only receive two samples of information: *X_pre_* represents the initial information and *X_post_* represents the additional (or post-decision) evidence. Both *X_pre_* and *X_post_* are sampled from normal distributions:

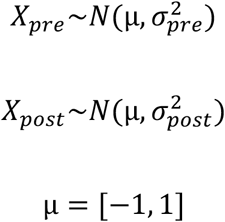

where the common mean (μ) of these distributions corresponds to the actual world state that needs to be inferred. While pre- and post-decision evidence distributions have the same mean (i.e. indicate the same underlying world state), they might differ in their variance. The variance represents the reliability of the information, with higher variances indicating less reliable information. We simulate different reliabilities of pre- and post-decision evidence as the consequences of a confirmation bias are likely to depend on this balance.

After receiving initial information, agents make an initial decision (*decision_initial_*) which depends solely on *X_pre_*. If X_pre_ has a positive value (*X_pre_*>0), *decision_initial_*=1, whereas if *X_pre_* has a negative value (X_pre_<0), *decision_initial_*=-1. An estimate of confidence in this initial decision is derived by calculating the log-odds in favour of the chosen world state:

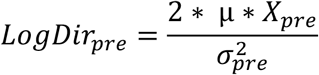

These log-odds can be transformed into a probability of being correct (between 0 and 1) as follows:

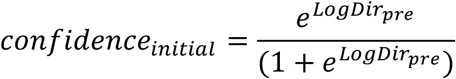

After the initial decision the agent receives additional information in the form of *X_post_*. In order to reach a final decision, both evidence samples can be integrated in an unbiased Bayesian fashion by simply summing the log-odds:

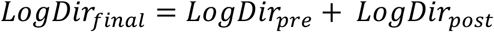

Note that by summing the log-odds of pre- and post-decision evidence, the certainty/reliability of these two evidence samples is implicitly considered, i.e. evidence is combined in line with Bayesian principles. The final decision depends on the sign of the posterior log-odds (*LogDir_final_*). If the final decision corresponds to the actual state of the world, the agent can be said to have formed an accurate belief and performs the task correctly. We refer to this unbiased decision-maker as a Bayesian agent.

A confirmation bias can be modelled within this framework as an altered incorporation of post-decision evidence dependent on whether this new information confirms or disconfirms the initial decision:

If sign(*X_post_*) = sign (*decision_initial_*):

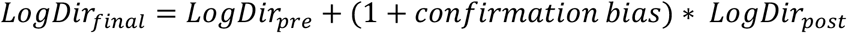

Else if sign(*X_post_*) ≠ sign (*decision_initial_*):

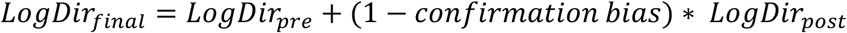

This form of confirmation bias can range from 0 (no bias) to 1 (where processing of disconfirmatory information is abolished).

In line with empirical observations, we also modelled a situation in which confirmation bias is modulated by confidence (Rollwage et al., 2020), such that participants were relatively unbiased in their use of new evidence when less confident, but showed an enhanced confirmation bias after high confidence decisions. To mimic these signatures, we simulated a “metacognitive” agent which shows a confirmation bias when it is confident in an initial choice (confidence=1), but remains unbiased when unsure (confidence=.5):

If sign(*X_post_*) = sign (*decision_initial_*):

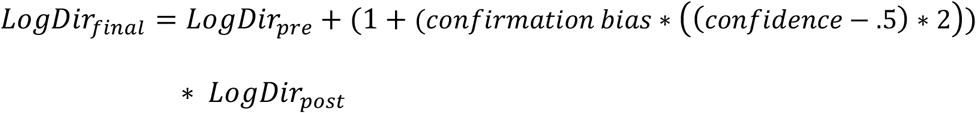

Else if sign(*X_post_*) ≠ sign (*decision_initial_*):

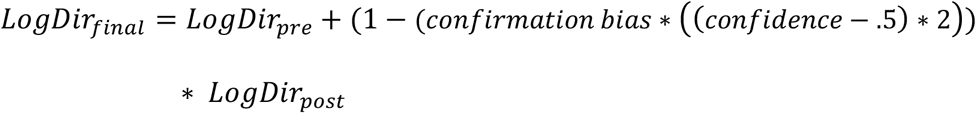

We note that a metacognitive agent differs from a Bayesian agent in that its initial confidence directly modulates the extent to which post-decision evidence is incorporated. For a Bayesian agent there is of course also a sense in which confidence “weights” the subsequent incorporation of evidence, in that a highly confident decision will require more disconfirming evidence to be overturned. Such updates, however, are in keeping with the linear accumulation of the log-odds of one or other hypothesis. In contrast, our metacognitive agent downweights the processing of disconfirmatory evidence when it is confident, representing a non-linear effect of confidence on the incorporation of post-decisional log-odds. Thus, our metacognitive agent shows similarities with circular inference models (Bouttier et al., 2019; Jardri et al., 2017), in which prior beliefs directly alter sensitivity to new evidence.

For each agent and each combination of evidence strength (see below) we simulated 200,000 trials, with the average decision accuracy across these simulated trials forming our measure of the agent’s performance. In what follows we describe in detail how these simulations were conducted.

## Methods

### Pre- and post-decision evidence strength

As the performance of the different agents might depend on the reliabilities of pre- (*X_pre_*) and post-decisional (*X_post_*) information, we simulated different information strengths defined as z-scores 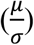. We fixed *μ*=1 (or −1 respectively) and changed σ_pre_ and σ_post_ to vary information strength. We modelled all combinations of pre-decision 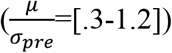 and post-decision 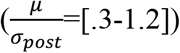 information strengths. In Figures 1A, B and D analyses of the joint effects of these two evidence strengths are presented, whereas in Figures 1C and 2A, C performance is averaged over all evidence strength conditions.

**Figure 1.**
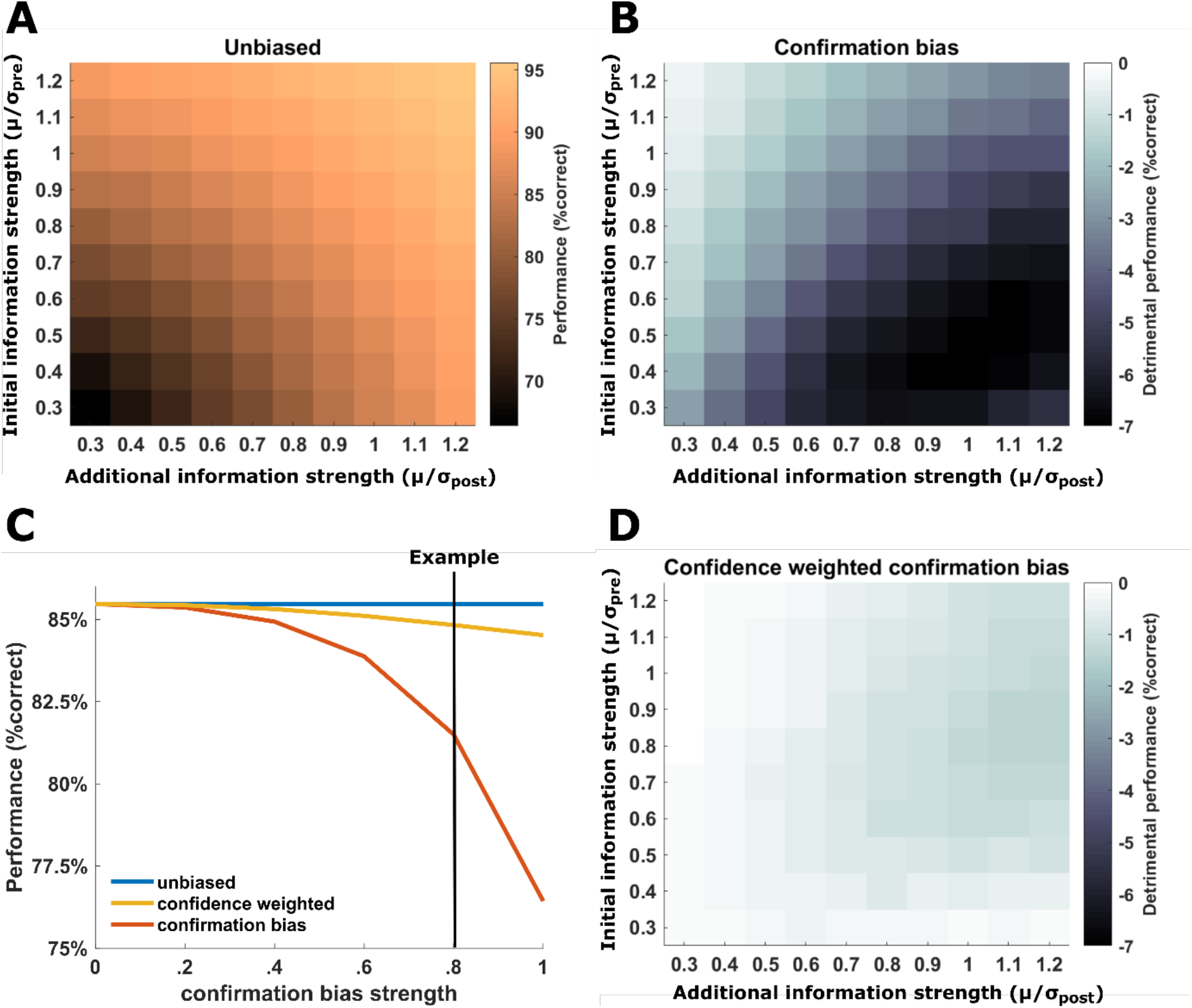
Comparison of agents’ performance with different biases in information processing. **A** Performance of an unbiased agent that integrates initial and additional information in a Bayesian manner. Depending on the evidence strength, this agent shows different levels of accuracy, with better performance when both evidence samples are strong/reliable. **B** Difference in performance between an unbiased agent and a confirmation-biased agent as a function of the reliability of the initial and additional evidence. A confirmation bias has especially detrimental effects when initial evidence is relatively weak. **C** Comparison of unbiased, confirmation-biased and metacognitive (confidence-weighted) agents as a function of confirmation bias strength. Performance is averaged over all combinations of initial and additional evidence strengths. The vertical line indicates the strength of confirmation bias used in panels B and D. **D** Difference in performance between an unbiased agent and a metacognitive agent as a function of the reliability of initial and additional evidence. Overall, the metacognitive agent shows only a relatively small disadvantage in comparison to an unbiased agent. In comparison to a simple confirmation bias, the metacognitive agent suffers less performance detriment in situations with weak initial evidence. **B&D** Dark colours indicate more detrimental performance of confirmation bias strategies when compared to unbiased evidence integration.

**Figure 2.**
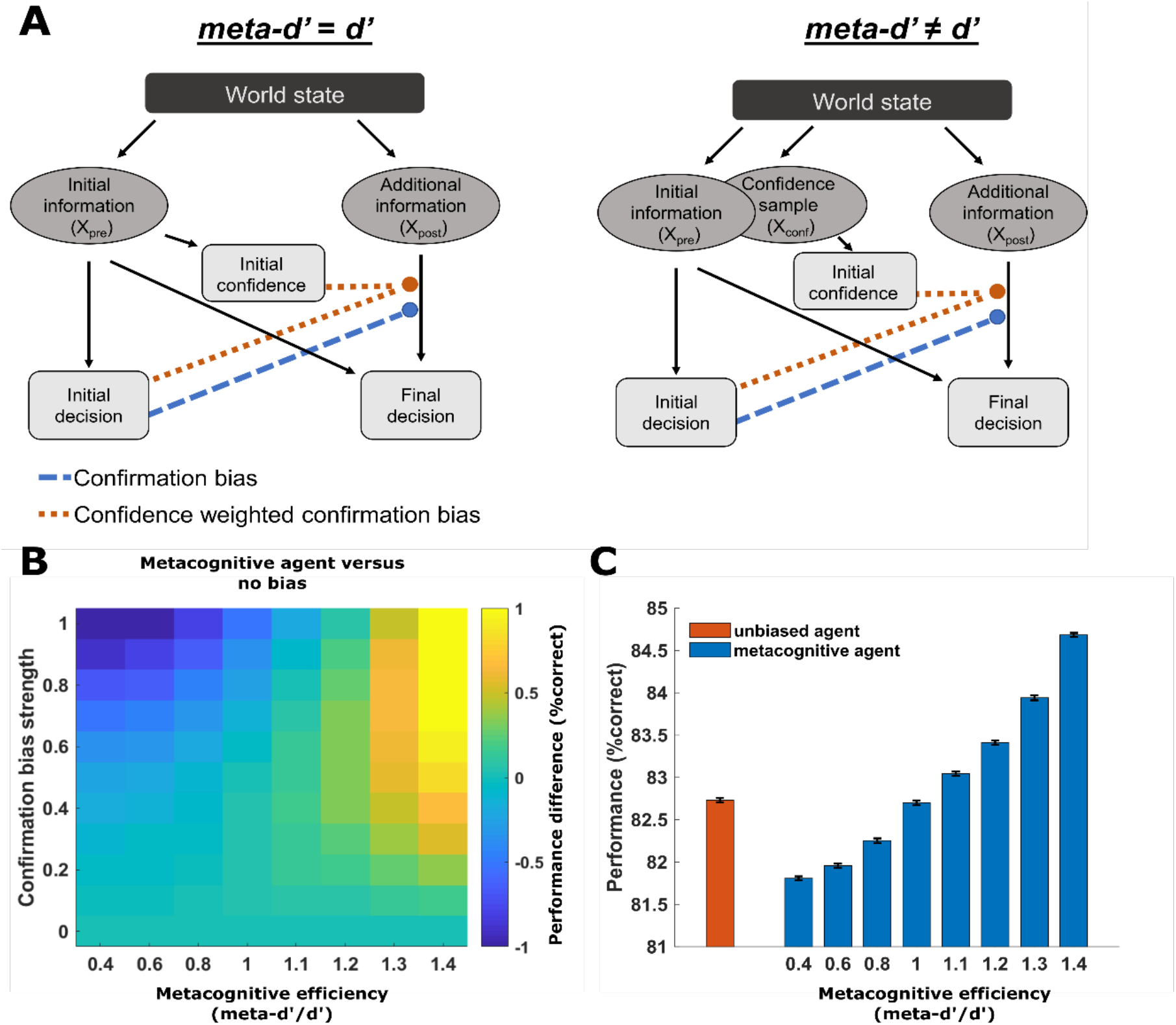
Performance of a metacognitive agent as function of metacognitive efficiency. **A** Illustration of different decision and confidence models. The left panel shows a model in which the same evidence (X_pre_) informs both the initial decision and the initial confidence, resulting in a ratio of meta-d’/d’=1. The right panel shows a situation in which meta-d’ and d’ can differ as the initial decision and confidence rely on separate, though correlated, evidence samples (X_pre_ & X_conf_). The final decision is determined by a combination of the initial (X_pre_) and the additional (X_post_) information. The coloured arrows indicate the way in which either an initial decision (confirmation bias) or an initial decision in combination with confidence (confidence-weighted confirmation bias, as in our metacognitive agent) modulate the incorporation of post-decision evidence. **B** Performance difference for a metacognitive agent compared to an unbiased agent. The performance of a confidence-weighted confirmation bias is sensitive to metacognitive efficiency (i.e. the accuracy of confidence ratings), with the greatest benefits obtained when metacognitive ability is high. When the ratio of meta-d’/d’ is above 1, a metacognitive agent outperforms an unbiased agent. Hotter colours indicate better performance of the metacognitive agent. **C** Performance of an unbiased agent compared to metacognitive agents differing in their metacognitive efficiency. Here we fix the pre- and post-decision evidence strengths to an intermediate level (see Methods) in order to reveal differences between different decision strategies. We simulated 100 agents for each setting and present group means ± SEM. Metacognitive agents with lower metacognitive efficiencies (meta-d’/d’<1) show significantly lower performance than an unbiased agent (all p<.0001), whereas metacognitive agents with higher metacognitive efficiencies (meta-d’/d’>1) show significantly better performance than an unbiased agent (p<.0001). An agent with meta-d’/d’=1 yields performance that is not significantly different from that of an unbiased agent (p=.32).

### Generative model of metacognitive abilities

The correspondence between confidence and performance can be formally quantified as the ratio of *meta-d’/d’* (known as metacognitive efficiency), where *meta-d’* reflects metacognitive sensitivity and *d’* reflects primary task performance within a signal detection theoretic framework (Maniscalco & Lau, 2012). Several reasons for a dissociation between *meta-d’* and *d’* have been suggested. For instance, confidence may reflect a noisy read-out of the decision evidence or a decline of decision evidence in working memory prior to a confidence judgment (Maniscalco & Lau, 2012), leading to a *meta-d’/d’* ratio of less than 1. On the other hand, confidence might be informed by evidence that was not available at the time of decision (Moran et al., 2015; Moreira et al., 2018; Pleskac & Busemeyer, 2010), or on correlated evidence that is accumulated in parallel (Fleming & Daw, 2017), both of which may lead *meta-d’/d’* ratios to surpass 1.

To model different degrees of metacognitive ability, we relaxed our assumption that confidence is directly derived from the evidence informing the initial decision (see Figure 2A). Instead, the evidence informing decisions (*X_pre_*) and confidence estimates (*X_conf_*) were modelled as distinct but correlated, and we allowed the reliability of the confidence 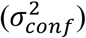 and decision 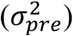 samples to differ. Specifically, *X_pre_* and *X_conf_* were sampled from a bivariate normal distribution with mean *μ* and covariance *Σ*:

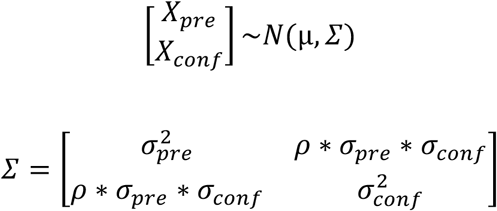

*Σ* parameterises the relationship between *X_pre_* and *X_conf_*, with *ρ* representing the correlation between these variables. As described by Fleming & Daw (2017), in this situation confidence can be inferred based on a combination of *X_conf_*, the initial decision, and the covariance of *X_conf_* and *X_pre_* (see the Appendix of Fleming & Daw, 2017, for further details on this calculation):

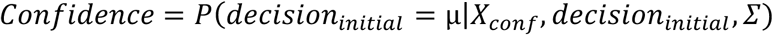

Importantly, modelling separate samples of X_conf_ and X_pre_ allows for dissociations between *meta-d’* and *d’*, thus making it possible to simulate varying degrees of metacognitive efficiency. For instance, a decrease in the reliability of the evidence informing the confidence rating 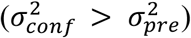 naturally reduces metacognitive efficiency and leads confidence judgments to be less reliable predictors of choice accuracy. For simulating lower metacognitive abilities (*meta-d’/d’*<1) we fixed *ρ*=.8 and varied 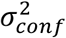 (however, we note that our findings are not dependent on the specific value of *ρ* or 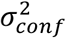; see Figure S1). 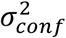 was defined by multiplying 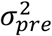 by the set of coefficients [2.04, 1.56, 1.23], which were selected to result in ratios of *meta-d’/d’* of [.4 .6 .8]. We also modelled a situation in which confidence and decision information have the same reliability 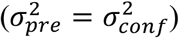, but the evidence samples have a variable degree of correlation (*ρ*=[.1, .3, .5, .65]), resulting in values of *meta-d’/d’* of [1.4, 1.3, 1.2, 1.1]. Such decorrelations in evidence samples result in increased metacognitive efficiency because there is additional information on which to base an evaluation of the decider (Fleming & Daw, 2017).

### Assessment of metacognitive efficiency

To assess whether manipulations in our generative model of confidence had the intended influence on agents’ metacognitive efficiency, we calculated the *meta-d’*/*d’* ratio for each agent (Maniscalco & Lau, 2012) based on their simulated behaviour, using the MLE toolbox of Maniscalco and Lau (http://www.columbia.edu/~bsm2105/type2sdt/). Model fits were conducted separately for each level of evidence strength and then averaged over all evidence strengths for each agent.

### Robustness and significance

We simulated many trials per condition (200,000 trials for each combination of pre- and post-decision evidence strength) to ensure robustness to noise perturbations. We also sought to provide a statistical test of the modulation of performance by metacognitive efficiency. We simulated 100 agents per metacognitive efficiency setting (with confirmation bias=1) and compared their performance to an unbiased agent (also simulated 100 times) using a t-test. We also evaluated the magnitude of effects of metacognition on post-decision performance at intermediate pre-decision evidence 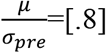 and post-decision evidence 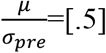 strengths, which are similar to those commonly used in previous laboratory experiments.

## Results

Depending on the different reliabilities of *X_pre_* and *X_post_*, an unbiased Bayesian observer achieves different final performances (see Figure 1A), with more reliable information yielding better performance. In addition to an unbiased observer, we also modelled agents with gradually increasing levels of confirmation bias (see Figure 1C). As hypothesized, selectively accumulating confirmatory information results in detrimental performance, with higher levels of confirmation bias leading to a more pronounced detriment. At higher levels of confirmation bias, a detriment in performance from 85% correct to ~77% correct was observed, which is notable in a 2AFC choice scenario where performance can only vary between 50 and 100% and evidence strength is adjusted to be of intermediate difficulty. As expected, this effect was most noticeable when the agent received relatively weak initial information but reliable post-decision evidence (see Figure 1B), as the bias prevents the incorporation of more reliable corrective information.

We next examined the performance of a metacognitive agent with a confidence-weighted confirmation bias. Notably, a metacognitive agent outperforms a simple confirmation bias in all settings and only shows slight impairments in relation to an unbiased agent (see Figure 1C). We also found that a metacognitive agent clearly outperforms a simple confirmation bias in situations when the initial evidence is weak (see Figure 1D). While a simple confirmation bias has the strongest decrement in performance in these situations, a metacognitive agent “realizes” it is dealing with weak initial evidence (by assigning lower confidence to these decisions) and thus shows a more equal sensitivity to confirming and disconfirming information.

Up until now we have assumed that agents calculate confidence in an initial choice by directly evaluating the reliability of evidence that informed the decision, resulting in a fixed metacognitive efficiency (*meta-d’/d’*=1). However, empirical evidence shows that people differ in their metacognitive ability (Ais et al., 2016; Fleming et al., 2010; Rollwage et al., 2018). Thus, we next assessed the degree to which metacognitive efficiency influences the performance of our metacognitive agent.

We found that reduced metacognitive efficiency (*meta-d’/d’*<1) leads metacognitive agents to show impaired performance compared to unbiased agents (all p-values<.0001, see cooler colours in Figure 2B), as the weighting of new information by confidence becomes less effective. Conversely, when metacognitive efficiency is high (*meta-d’/d’*>1) we found that a metacognitive agent can even outperform an unbiased agent (all p-values<.0001, see hotter colours in Figure 2B). This result is striking as it suggests that a selective integration of information is not necessarily a “bias” but represents an advantageous strategy for achieving optimal performance in the context of a realistic cognitive architecture (i.e. one in which metacognition is particularly efficient). Figure 2C displays the effect of metacognitive efficiency on the performance of a metacognitive agent at a fixed level of intermediate evidence strength, relative to an unbiased agent. While the magnitude of these differences might appear relatively small (of the order of 2-3% correct), individual differences between human participants performing a similar task cover a similar range (e.g. in Rollwage et al. 2020 the standard deviation of performance across participants was 3.7% correct decisions).

## Discussion

Here we investigated the effects of confirmation bias on the accuracy of belief formation. Our central proposal is that when confirmation bias is a feature of a self-aware (metacognitive) agent, it ceases to be detrimental, and may even become adaptive. We used simulation-based modelling to compare the performance of agents with different forms of confirmation bias against an unbiased agent. A simple (non-metacognitive) confirmation bias showed detrimental effects compared to an unbiased agent in all settings. In comparison, a metacognitive agent which modulates the degree of confirmation bias by confidence (as documented empirically in human observers; Rollwage et al., 2020) outperformed a simple confirmation bias agent, and was in many cases not substantially worse than an unbiased agent. The benefit of weighting a confirmation bias by confidence is that when confidence is low, and errors are more likely, the system becomes open to new and potentially corrective information.

In turn, by simulating varying degrees of metacognitive efficiency, we found that the performance of our metacognitive agent was sensitive to its level of self-awareness. Strikingly, a metacognitive agent with high self-awareness could in some cases even outperform an unbiased agent, indicating that selective information processing might be particularly adaptive when coupled with good metacognitive abilities. These results are in accordance with a view that cognitive biases may have originally evolved for good evolutionary reasons and are often adaptive when considered in the context of the agent’s environment, including its broader mental toolkit (Gigerenzer, 2008).

Why should high degrees of metacognitive efficiency be advantageous in this case? The core mechanism appears to be the capacity of a confidence estimate to provide a “second look” on a decision, similarly to how an external adviser might give us a separate view on a topic. The benefit of this mechanism depends on the agent’s metacognitive ability – as agents with good metacognition provide the most effective “internal” advisory signals. Interestingly, however, a confidence-weighted confirmation bias outperformed a simple confirmation bias in all settings, even when metacognitive ability was relatively low (see Figure S2), suggesting that the mere presence of confidence-weighting, rather than acute metacognition per se, may be sufficient to avoid the most deleterious effects of confirmation bias.

While these advantages for metacognitive agents were relatively small in size, they were robust and similar in magnitude to individual differences in performance on comparable laboratory tasks (e.g. Rollwage et al., 2020). We note that here we modelled a situation in which only two consecutive samples of evidence had to be integrated. Even in this minimal paradigm, increases in metacognitive efficiency could lead to a 2-3% increase in the number of correct decisions. Such a bias towards higher performance on individual, isolated judgments is likely to be magnified in situations requiring the integration of multiple information samples over time. In such situations, even small alterations in information processing might summate over time and lead to substantial changes in belief accuracy.

By incorporating confirmation bias as part of a broader cognitive architecture in which different mental processes can interact with each other (e.g. decisional and metacognitive processes), selective information processing may become adaptive when compared to the same “bias” considered in isolation. In the same spirit, it has been argued that heuristics that may appear as biases in simple and constrained laboratory tasks become beneficial in more complex environments (Pleskac & Hertwig, 2014). More broadly, our study indicates that considering cognitive biases in isolation from other mental processes might lead to the wrong conclusions about the impact of a particular cognitive feature on behaviour.

We note that a behavioural benefit for confirmation bias was only present when simulating agents with “hyper” metacognitive efficiency (i.e. *meta-d’/d*>1), such that metacognition became more acute than first-order task performance. This might seem odd at first glance, as it implies that the system is not using all the information available to it at the time of making an initial choice, and only afterwards becomes more sensitive to whether it was right or wrong. However, this kind of pattern is commonly observed in empirical data (Fleming & Daw, 2017; Moreira et al., 2018), and is thought to be driven either by additional post-decisional processing, differences in the variance of signal compared to noise (Miyoshi & Lau, 2020) or (as simulated here) parallel streams of information processing that allows the system to detect and correct its errors (Moreira et al., 2018). The capacity for rapid error detection is well established in human studies (Murphy et al., 2015; Rabbitt, 1966; Resulaj et al., 2009; Van Den Berg et al., 2016; Yeung & Summerfield, 2012) and thus it is reasonable to assume that hyper metacognitive efficiency may be common in the healthy population.

Importantly, a metacognitive system does not need to be more reliable than the decision-maker to achieve high metacognitive ability: it needs only to incorporate partially independent information (as used here; see Figure S1 for simulations using a wider set of parameters). In this respect, our results also contribute to a debate over why it might be useful for the brain to encode a confidence signal separately from representations of decision evidence (Maniscalco et al., 2016; Meyniel et al., 2015). A metacognitive agent that can realize its own mistakes (and assign low confidence to these decision) will tend to become more open to new information due to the confidence weighting applied to selective information processing (interestingly our model predicts that when confidence falls below 0.5 in a two-choice scenario, metacognitive agents should even show a “disconfirmation” bias, and be more prone to seek out information contradicting their current position). Our results suggest that this benefit is only accrued when metacognition is partly independent of first-order cognition.

Selective information processing has been assumed to lead to skewed, entrenched and potentially inaccurate beliefs about a range of societal and political issues (Rollwage et al., 2019; Zmigrod, 2020). However, the current results suggest that the detrimental effects of selective information processing depend on people’s broader self-awareness. In turn, metacognitive deficits might represent core drivers of polarised or radical beliefs, due to their consequence for maladaptive confirmation bias. Interestingly, this hypothesis is in line with empirical observations (Rollwage et al., 2018), showing that more dogmatic participants show reduced metacognitive sensitivity which in turn is predictive of reduced post-decision evidence processing.

Recognising metacognition as a central driver of belief polarisation may make it possible to develop new strategies for debiasing decision-making (Rollwage et al., 2019). The contributors to confirmation bias in any given setting are likely to be multifactorial, with more proximal causes (such as measures of cognitive style) having large effect sizes, but providing more limited mechanistic insight. In contrast, identifying small, reliable effect sizes associated with underlying mechanisms (such as the impact of confidence weighting) may bring us closer to the potential for targeted intervention. Excitingly, there are existing interventions that have been shown to boost people’s metacognitive ability (Baird et al., 2014; Carpenter et al., 2019). Our results indicate that cognitive training which improves domain-general self-awareness and metacognitive efficiency may help to alleviate the negative behavioural outcomes of selective information processing, and foster resilience against misinformation and belief polarisation.

## Acknowledgements

We thank Prof. Ralph Hertwig for insightful conversations regarding the potential adaptiveness of cognitive biases. M.R. is a predoctoral Fellow of the International Max Planck Research School on Computational Methods in Psychiatry and Ageing Research. The participating institutions are the Max Planck Institute for Human Development and University College London (UCL). The Wellcome Centre for Human Neuroimaging is supported by core funding from the Wellcome Trust (203147/Z/16/Z). S.M.F. is supported by a Sir Henry Dale Fellowship jointly funded by the Wellcome Trust and the Royal Society (206648/Z/17/Z).

## Author Contributions

M.R. and S.M.F. conceived the study and wrote the paper. M.R. developed the methodology and ran the simulations with input from S.M.F.

## Code availability

Code of all simulations for this study is available at a dedicated Github repository (https://github.com/MaxRollwage/ConfirmationBias_simulation)

## Supplementary material

In the main text we focussed on the influence of metacognitive efficiency on the performance of a confidence-weighted confirmation bias, showing that this information processing strategy can outperform an unbiased agent when metacognitive efficiency is high. However, when sampling *X_pre_* and *X_conf_* from separate but correlated distributions, metacognitive efficiency can be influenced by either *ρ, σ*_conf_ or both. Within such a model, metacognitive abilities are known to increase when the correlation (*ρ*) between X_pre_ and X_conf_ is reduced, as well as when X_conf_ is more reliable, i.e. σ_∞_nf is small (Fleming & Daw, 2017; see Figure S1A). Here we show that independently of how these boosts in metacognitive efficiency are achieved, the results we obtain on the benefits for post-decision processing are maintained (Figure S1B). In other words, the performance of a metacognitive agent compared to an unbiased agent is determined by its metacognitive efficiency, but not the particular parameters of the confidence model used to generate variation in metacognitive efficiency. This inference is supported by a very high correlation between variation in metacognitive efficiency and post-decision integration performance when both are computed as a function of both *ρ* and *σ*_conf_ (r=.97, Figure S1C).

**Figure S1.**
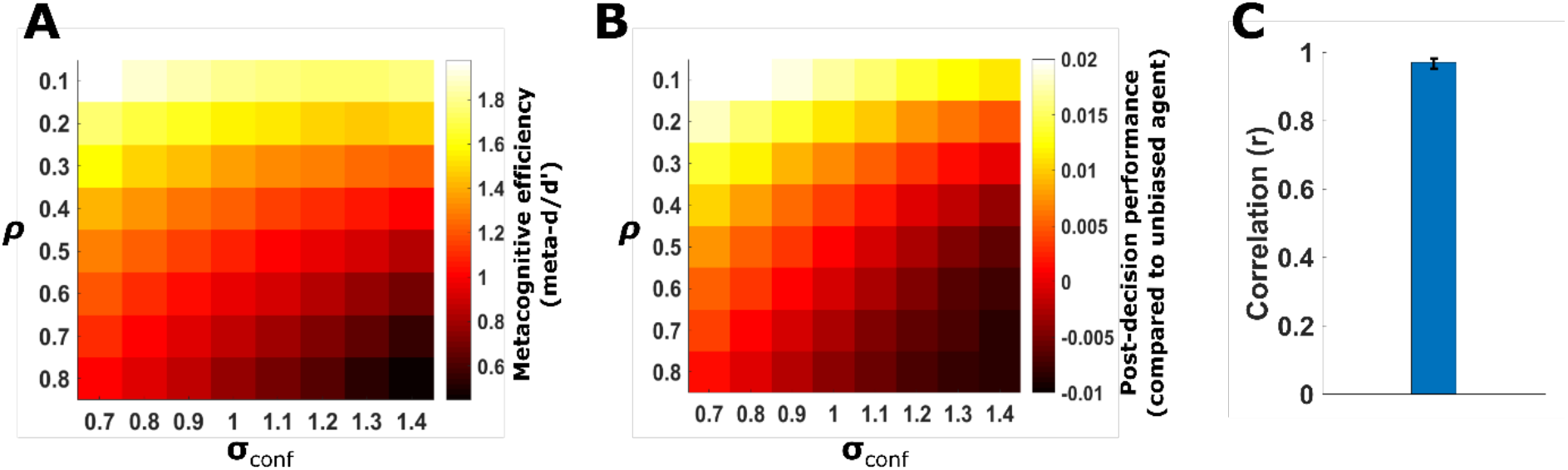
Relationship between metacognitive efficiency and post-decision integration performance. **A** The agent’s metacognitive efficiency (meta-d’/d’) is determined by the settings of σ_conf_ and ρ. Lower values of σ_conf_ indicate that confidence relies on more reliable evidence samples, whereby lower values ofρ mean that the evidence for the decision and confidence are only weakly correlated, i. e. both provide independent information. **B** Performance difference of a metacognitive agent compared to an unbiased agent, as a function of σ_conf_ and ρ. These parameters have very similar influences on both metacognitive efficiency and post-decision evidence integration (compare panels A and B). This indicates that, independently of how a specific metacognitive ability is achieved, it has the same influence on post-decision evidence integration. **C** Support for a link between metacognition and post-decision processing benefit is obtained by computing the correlation between these statistics across all values of σ_conf_ and ρ, i.e. the correlation between panels A and B (± 95% confidence interval).

**Figure S2.**
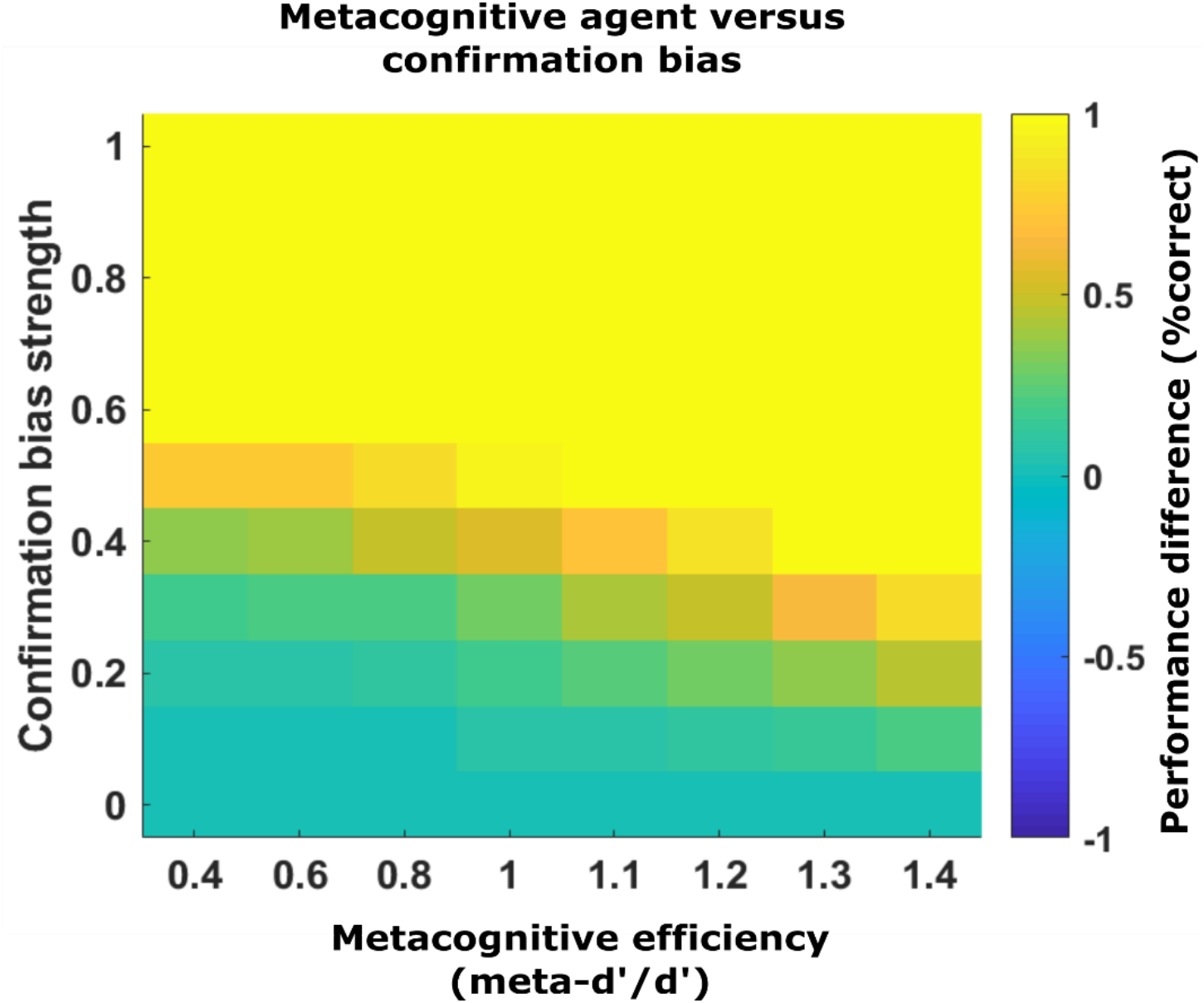
Performance difference of a metacognitive agent compared to a simple confirmation bias. Hotter colours indicate better performance of the metacognitive agent compared to a simple confirmation bias. The metacognitive agent shows superior performance in all settings, even when metacognitive efficiency is low.

